# The secreted elastase SjCE2b drives host skin penetration by *Schistosoma japonicum*

**DOI:** 10.64898/2026.07.12.738118

**Authors:** Bingkuan Zhu, Yi Shen, Fang Luo, Chuan Su, Hong You, Xumin Zhang, Wei Hu

**Affiliations:** State Key Laboratory of Genetics and Development of Complex Phenotypes, Ministry of Education Key Laboratory of Contemporary Anthropology, School of Life Sciences, Fudan University, Shanghai, China; Institute of Biomedical Sciences, College of Life Sciences, Inner Mongolia University, Hohhot, China; National Institute of Parasitic Diseases, Chinese Center for Disease Control and Prevention (Chinese Center for Tropical Diseases Research), NHC Key Laboratory of Parasite and Vector Biology, WHO Collaborating Center for Tropical Diseases, National Center for International Research on Tropical Diseases, Shanghai, China; School of Public Health, LKS Faculty of Medicine, The University of Hong Kong, Hong Kong SAR, China; Key Laboratory for Pathogen Infection and Control of Jiangsu Province, Department of Pathogen Biology and Immunology, School of Basic Medical Sciences, National Vaccine Innovation Platform, Nanjing Medical University, Nanjing, Jiangsu, China; Infection and Inflammation Program, QIMR Berghofer Medical Research Institute, Brisbane, QLD, Australia

**Keywords:** *Schistosoma japonicum*, cercariae, secretome, elastase, SjCE2b, skin penetration

## Abstract

Cercarial elastase is the most abundant protease secreted by *Schistosoma mansoni* and plays a critical role in cercarial invasion. Although *Schistosoma japonicum* encodes only a single elastase, SjCE2b, its secretion by cercariae and its specific function in skin penetration have remained elusive. Here, we report the first proteomic analysis of *S. japonicum* cercarial excretory-secretory products (ESPs) induced by linoleic acid or mouse skin, confirming the presence of SjCE2b in both ESPs preparations. Recombinant SjCE2b expressed in *Pichia pastoris* was characterized as a trypsin-like serine protease whose activity is entirely abolished by the elastase inhibitor MeoSuc-AAPF-CMK. Furthermore, SjCE2b expression was detected exclusively in cercarial extracts and localized specifically to the cercarial acetabular glands and ducts. Subsequent proteomic analysis indicates that SjCE2b can degrade numerous human epidermal proteins, including ten isoforms of type I and type II keratins. *In vitro* digestion assays further demonstrated that SjCE2b can digest key structural components of the dermis, including elastin, collogen, and fibronectin. Additionally, the cleavage of complement component C3 and immunoglobulins (IgA and IgG) suggests that SjCE2b may facilitate immune evasion by newly transformed schistosomula. Critically, the incubation of cercariae with anti-rSjCE2b antibody reduced the worm burden by 80.85%, confirming the essential role of SjCE2b in the skin penetration of *S. japonicum* cercariae and highlighting it as a compelling candidate for vaccine or therapeutic development.

**Author Summary:** Schistosomiasis constitutes a major global health burden caused by parasitic flatworms of the genus *Schistosoma*. Proteolytic and histolytic enzymes secreted by cercarial pre- and post-acetabular glands facilitate disruption of the host skin barrier and protect the parasite from localized dermal inflammatory response. Although cercarial elastase is a well-characterized invasion enzyme in *Schistosoma mansoni*, its role in *Schistosoma japonicum* remains poorly defined; consequently, *S. japonicum* cercariae have been hypothesized to rely on distinct repertoire of proteolytic enzymes during skin penetration. In the present study, we showed that SjCE2b was localized to the cercarial acetabular glands and ducts, and was secreted upon stimulation with linoleic acid or mouse skin. Functionally, SjCE2b can disrupt host skin integrity by degrading epidermal and dermal components, and may promote immune evasion through cleavage of complement component and immunoglobins. Furthermore, the antibody-mediated neutralization of secreted SjCE2b significantly impaired parasite penetration by greater than 80%. Together, these findings establish SjCE2b as a critical enzyme required for *S. japonicum* cercariae invasion and highlight its potential as a promising target for novel therapeutic interventions.

## Introduction

Schistosomes, or blood flukes, are the etiological agents of schistosomiasis. It is estimated that approximately 250 million people are infected with schistosomes across 78 countries, with around 1 billion individuals at risk of infection (1). Among the six clades of *Schistosoma* genus, *Schistosoma haematobium*, *Schistosoma mansoni* and *Schistosoma japonicum* are the primary agents of human diseases (2). The life cycle of schistosomes involve two hosts: a freshwater snail as the intermediate host and a mammal as the definitive host. Following intra-molluscan maturation, infective cercariae are shed into the surrounding water in response to light. Upon encountering a suitable host, they will actively penetrate the exposed skin using proteolytic and histolytic enzymes secreted by their acetabular glands (3). Notably, *S. japonicum* traverses the epidermis and dermis much faster than *S. mansoni* and *S. haematobium*, requiring 3 days, 5-7 days and >7 days, respectively (4). Thus, a different penetration mechanism is speculated to be employed by *S. japonicum*.

During definitive host invasion, *Schistosoma* cercariae release acetabular gland contents enriched with potent histolytic proteases and bioactive molecules that facilitate skin penetration and modulate the host immune response (5). While cercarial excretory-secretory products (ESPs) of *S. mansoni* have been characterized using different stimulation methods, including mechanical transformation by passage through a 22-gauge needle (6) or vortexing (7), as well as exposure to human skin lipid (8) or human skin (9). However, ESPs of *S. japonicum* have so far been collected and analyzed only after mechanical transformation by passage through a 22-gauge needle (6), a procedure that may damage parasite bodies and lead to contamination with somatic proteins.

The most abundant protease identified in *S. mansoni* cercarial secretions is the serine protease cercarial elastase (SmCE). While the *S. mansoni* genome encodes an expanded family of eight full-length elastase genes (SmCE1a.1, SmCE1a.2, SmCE1b, SmCE1c, SmCE2a.1, SmCE2a.2, SmCE2a.3, and SmCE2b) (10), and S. haematobium possesses four (ShCE1a, ShCE1b, ShCE2a, and ShCE2b) (11), the *S. japonicum* genome harbors only a single elastase gene, SjCE2b (12). SmCE facilitates host penetration by degrading major skin structural proteins—including elastin, collagen, keratin, and fibronectin—and promotes immune evasion by cleaving complement component C3 (13, 14). Inhibition of serine protease activity with chemical inhibitor reduces *S. mansoni* cercarial invasion by 97–100%, highlighting the essential role of SmCE in cercarial penetration (15). Despite these insights, the functional biology of the *S. japonicum* homolog remains largely uncharacterized. Specifically, the expression profiles of SjCE2b during larval development, whether it is secreted during host penetration, and its potential roles in tissue degradation and immune evasion represent critical knowledge gaps that warrant comprehensive investigation.

In the present study, we performed proteomic analysis of *S. japonicum* cercarial ESPs following stimulation with linoleic acid, a fatty acid known to induce ESP release and tail shedding in cercariae (16), as well as exposure to host skin, which more accurately simulates the natural penetration process. SjCE2b was identified in the secretome under both conditions. We further demonstrated that SjCE2b is expressed in cercariae and localized to the acetabular glands and gland ducts. In addition, we purified the functional enzyme and found that it can induce substantial degradation of both the epidermal and dermal layers of host skin. Collectively, these findings expand current knowledge of the *S. japonicum* cercarial secretome and reveal a critical functional role for SjCE2b during the host tissue invasion.

## Materials and Methods

### Ethics statement

All animal experiments were conducted in accordance with the Guidelines for the Care and Use of Laboratory Animals of the Ministry of Science and Technology of the People’s Republic of China (2006398) and were approved by the Ethics and Animal Welfare Committee of the National Institute of Parasitic Diseases, Chinese Center for Disease Control and Prevention, Shanghai, China (IPD2008-4).

### Parasites and animals

*S. japonicum* cercariae were isolated from Anhui province, China, and infected *Oncomelania hupensis* snails were provided by the Pathogen Biology Laboratory of the National Institute of Parasitic Diseases, Chinese Center for Diseases Control and Prevention, Shanghai, China. Six-week-old female C57BL/6 strain mice were purchased from Shanghai Animal Center, Chinese Academy of Sciences, Shanghai, China.

### Collection of *Schistosoma japonicum* cercarial secretions

To collect the excretory-secretory products (ESPs) of *Schistosoma japonicum* cercariae, approximately 40,000 cercariae floated in water were stimulated by either the introduction of a mouse tail or the addition of linoleic acid to a final concentration of 10 μL/mL. Following a 1-hour incubation, the suspension containing ESPs, live cercariae, and debris was chilled on ice and subjected to repeated centrifugation (13,000 rpm at 4 °C) to remove parasite bodies. Forty milliliters of the resulting supernatant were concentrated across six consecutive 15-minute centrifugation cycles (5,000 × g, 4 °C) using an Amicon Ultra-15 centrifugal filter unit with a 10 kDa molecular weight cutoff (Millipore). Total protein yield was quantified via Bradford assay (Pierce Biotechnology), and the final samples were lyophilization and stored for subsequent analysis.

### Mass spectrometry and ESPs protein identification

Enzymatic digestion of ESPs was performed using the filter-aided sample preparation (FASP) method as previously described (17). Briefly, 100 μg of protein per sample was reduced with 10 mM DTT at 56 °C for 45 min and subsequent alkylated with 100 mM iodoacetamide (IAM) in the dark at room temperature for 45 min with constant shaking. Samples were transferred to 10 kDa molecular weight cutoff filters (VWR International), adjusted to 400 μL with a buffer containing 100 mM Tris-HCl, 8 M urea, and 50 mM TEAB (pH 7.5), and centrifuged at 13,800 × g for 60 min. The retained protein pellets were washed once with 400 μL of 100 mM Tris-HCl and 50 mM TEAB (pH 7.5), followed by a final wash with 400 μL of 50 mM TEAB (pH 7.5), each with centrifugation at 13,800 × g for 60 min. Proteins were digested by adding 4 μg of trypsin (Sigma-Aldrich) in 100 μL of 50 mM TEAB (pH 7.5) directly to the filter and incubating the assembly at 37 °C for 12 h. Resulting peptides were desalted using a C18 solid-phase extraction column (CNW) according to the manufacturer’s instructions.

For the linolenic acid stimulated ESPs samples, LC-MS/MS analysis was performed on a nanoflow EASY-nLC 1200 system coupled to an Orbitrap Exploris 480 mass spectrometer (Thermo Fisher Scientific). Samples were analyzed utilizing a single-column setup with an in-house packed C18 analytical column (75 µm i.d. × 20 cm, ReproSil-Pur 120 C18-AQ, 1.9 µm; Dr. Maisch GmbH) (18). The mobile phases consisted of 0.1% formic acid in water (Buffer A) and 0.1% formic acid in 80% acetonitrile (Buffer B). Peptides were eluted at a flow rate of 200 nL/min using the following gradient: 5–8% B for 2 min, 8–44% B for 38 min, 44–70% B for 8 min, 70– 100% B for 2 min, and a final hold at 100% B for 10 min. The mass spectrometer was operated in data-dependent acquisition (DDA) mode with a 1 s cycle time. Precursors were fragmented in HCD mode with a normalized collision energy (NCE) of 30.

For the mouse skin stimulated ESPs samples, LC-MS/MS analysis was performed on a nanoflow EASY-nLC 1200 system coupled to an Orbitrap Fusion Lumos mass spectrometer (Thermo Fisher Scientific). The mobile phases consisted of 0.1% formic acid in water (Buffer A) and 0.1% formic acid in 80% acetonitrile (Buffer B). Peptides were eluted at a flow rate of 200 nL/min using the following gradient: 2–5% B for 3 min, 5–35% B for 40 min, 35–44% B for 5 min, 44–100% B for 2 min, and a final hold at 100% B for 10 min. The mass spectrometer was operated in data-dependent acquisition (DDA) mode with a 2 s cycle time. Precursors were fragmented in HCD mode with a normalized collision energy (NCE) of 30.

For protein identification, raw data were searched against our SjV3 protein database (19) using Proteome Discoverer (version 1.4.0.288; Thermo Fisher Scientific) integrating the Mascot search engine (version 2.7.0; Matrix Science). Precursor and fragment ion mass tolerances were set to 10 ppm and 0.05 Da, respectively, allowing up to two missed cleavages. Carbamidomethylation of cysteine was specified as a fixed modification, whereas protein N-terminal acetylation and methionine oxidation were set as variable modifications. The false discovery rate (FDR) threshold was strictly set to 1%. Environmental and mouse skin contaminants were filtered from the results. Finally, all identified *S. japonicum* proteins were searched against the MEROPS 12.5 database (20) using BLASTp (E-value < 10⁻⁵) to identify potential proteases.

### Sequence analysis and molecular modeling of SjCE2b

The amino acid sequence of SjCE2b was retrieved from NCBI with the accession number ACR27083.1. Domain architecture and functional motifs were analyzed by SMART tool (http://smart.embl-heidelberg.de/smart/set_mode.cgi?NORMAL=1). A spatial model of SjCE2b was constructed via homology modeling, as previously described (21), utilizing the X-ray crystal structure of bovine chymotrypsin (PDB ID: 4CHA) as the structural template. The initial homology model was generated in MODELLER using the Basic Modeling method, followed by structural refinement via the Chiron web server (https://dokhlab.med.psu.edu/chiron/login.php) and SAVES v6.0 (https://saves.mbi.ucla.edu/). The *in silicon* molecular docking between SjCE2b and inhibitor Suc-AAPF-CMK was implemented by the AutoDock 4.2.6 software. Molecular images were generated with Pymol software.

### Expression of recombinant SjCE2b in *P. pastoris* and antibody preparation

Condon-optimized SjCE2b gene sequence was synthesized by GenScript and constructed into the expression vector pPIC9k. The plasmid SjCE2b-pPIC9k, was linearized using *Sal*I and transformed into *P. pastoris* strain GS115 via electroporation at 2 kV, 25 μF, 200 Ω in electroporation cuvettes (Bio-Rad). Geneticin (G418)-resistant *P. pastoris* transformants were initially screened on MD plates, followed by selection on YPD plates containing increasing concentrations of G418 (1.0, 2.0, and 4.0 mg/mL). Genomic integration of SjCE2b was confirmed by PCR using 5’-AOX1 and 3’-AOX1 primers, with template genomic DNA extracted via a Yeast Genomic DNA Extraction Kit (Thermo Fisher). Positive clones were inoculated into 100 mL of BMGY medium and cultured at 28 °C with constant shaking for 24 h. For protein induction, cells were harvested by centrifugation at 250 × g for 10 min at room temperature and resuspended in 1 L of BMMY medium. The culture was maintained for four days, with methanol added every 24 h to a final concentration of 1% (v/v). Post-induction, cells were pelleted at 2,500 × g for 10 min. The supernatant containing rSjCE2b was dialyzed against three volumes of PBS and purified via Ni^2+^-NTA affinity chromatography (GE Healthcare). The molecular weight and purity of rSjCE2b were assessed using 12% SDS-PAGE (GenScript) followed by Coomassie blue staining. Protein concentration was determined using a Bradford assay (Bio-Rad). Purified proteins were aliquoted and stored in PBS at -80 °C. The purification process was monitored via SDS-PAGE, Western blotting, and a kinetic activity assay utilizing the fluorogenic substrate Suc-AAPF-AMC (Sigma-Aldrich).

Polyclonal specific antibodies to purified recombinant SjCE2b were generated in rabbits (Shanghai Youke Biotechnology Co., Ltd.). Primary immunization was administered intraperitoneally on day 1 using antigen emulsified in Freund’s Complete Adjuvant. Subsequent booster immunizations formulated with Freund’s Incomplete Adjuvant were administered on days 15 and 43, while a negative control rabbit was mock-immunized with PBS alone. Immune serum was collected on day 53, and specific IgG fractions were isolated via affinity chromatography using a HiTrap Protein A column (GE Healthcare) according to the manufacturer’s protocol.

### RNA isolation, cDNA synthesis, and RT-qPCR

Eggs, miracidia, daughter sporocysts and cercariae were collected, washed three times with 1 mL PBS, resuspended in 500 μL TRIzol reagent (Takara), and processed for total RNA extraction as previously described (22). Total RNA was subsequently reverse-transcribed into cDNA using the PrimeScript RT Reagent Kit (Takara) according to the manufacturer’s instructions.

Transcriptional profiling of SjCE2b was conducted via RT-qPCR using specific primers (forward: 5’-AGATGGTCAACTTAGAGA-3’; reverse: 5’-CATATCGTCCTGATGTATC-3’). Amplification was performed in 10 μL reactions utilizing SYBR Green Master Mix (Yeasen) in 96-well plates (Genebrick). The thermal cycling conditions consisted of an initial denaturation at 95 °C for 6 min, followed by 40 cycles of 95 °C for 10 s and 60 °C for 20 s, concluding with a melt curve analysis (95 °C for 10 s, 65 °C for 60 s, 97 °C for 1 s) to confirm specific amplification. Samples were run in technical duplicates. Relative SjCE2b mRNA expression levels were normalized against the internal control PSMD (26S proteasome non-ATPase) and calculated using the 2^−ΔCt^ method (23).

### Western blot analysis

Ten μg of proteins extracted from eggs, miracidia, daughter sporocysts and cercariae were separated by SDS-PAGE and electrotransferred onto a PVDF membrane (Pall Corporation). Non-specific binding sites were blocked in 5% skim milk in TBS (pH 8.0) for 2 h at 37 °C. Following blocking, membranes were incubated overnight at 4 °C with either the primary anti-rSjCE2b IgG (diluted 1:2,000) or an anti-GAPDH antibody (diluted 1:2,000; Proteintech Group). After three 15-min washes in TBST, membrane was incubated with a horseradish peroxidase (HRP)-conjugated goat anti-rabbit IgG secondary antibody (diluted 1:5,000; Beyotime Biotechnology) for 2 h at room temperature. Following a final series of TBST washes, immunoreactive bands were detected using NcmECL Ultra solution (NCM Biotech) and imaged with a Tanon 5200 system (Tanon).

### Immunolocalization of SjCE2b in cercariae

Immunofluorescence analysis was performed as previously described (24). Samples were incubated with either the primary anti-rSjCE2b antibody (1 μg/mL) or a pre-immune IgG control (1 μg/mL). Another negative control, in which the primary antibody was omitted and samples were incubated in blocking solution alone, was also included.

### Protease activity assay

Proteolytic activity was measured using the fluorogenic peptide substrate Suc-AAPF-AMC (Sigma-Aldrich), which liberates the free fluorophore 7-amino-4-methylcoumarin (AMC) upon cleavage. Assays were performed in triplicate in black, flat-bottom 96-well microplates (Thermo Fisher) within a total reaction volume of 200 μL. Varying concentrations of recombinant SjCE2b (25, 50, and 100 nM) were pre-incubated for 10 min at 37 °C in 180 μL of standard assay buffer (50 mM Tris-HCl, pH 8.0). Enzymatic cleavage was initiated by the addition of 20 μL of the substrate solution (5 μM final concentration). Fluorogenic product release was monitored continuously for 30 min using a microplate reader (BioTek) at excitation and emission wavelengths of 380 and 460 nm, respectively. For inhibition assays, specific protease inhibitors (at the final concentrations listed in Table 1) were introduced during the 10-min enzyme pre-incubation step prior to substrate addition.

**Table 1.**
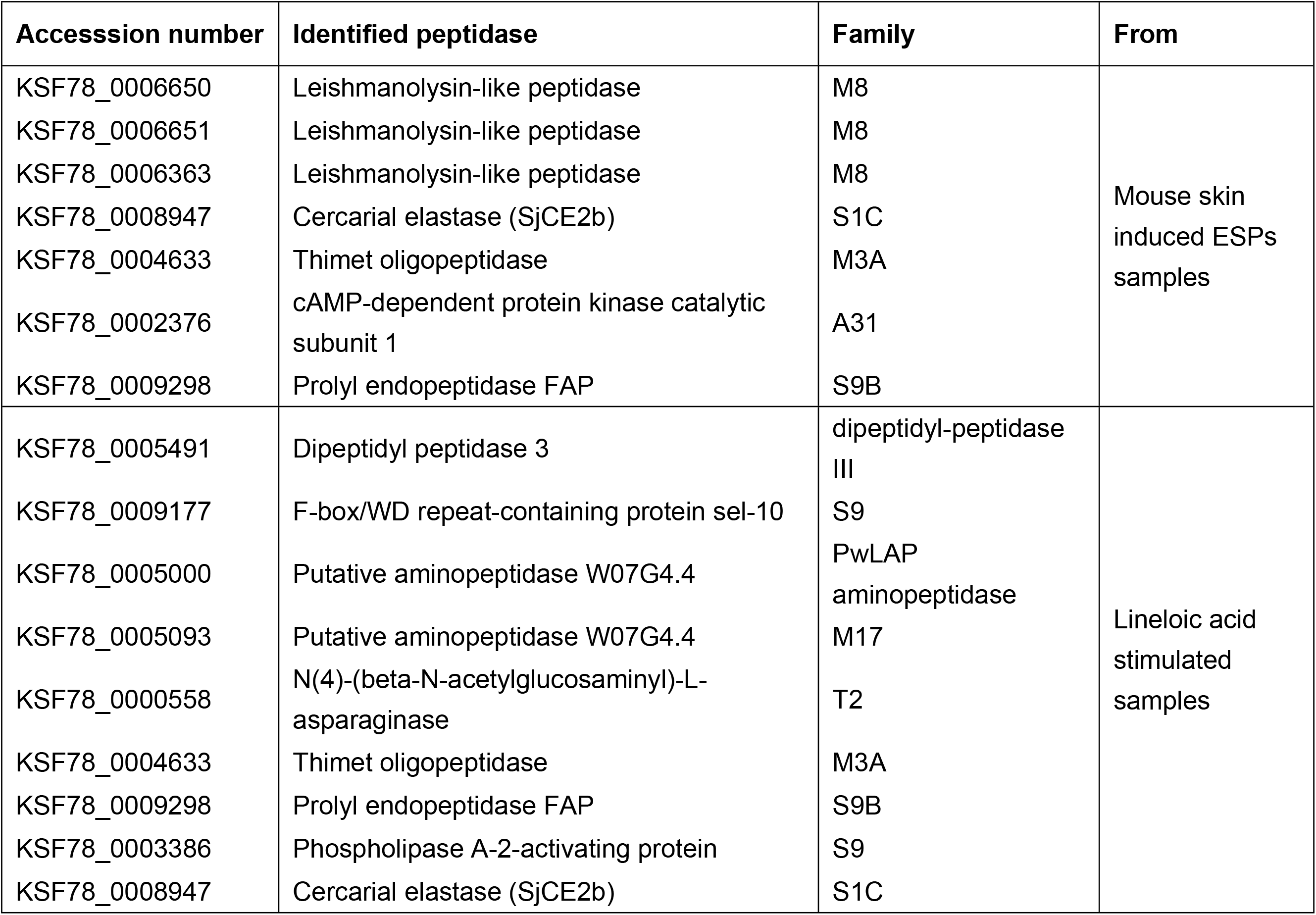
Peptidases identified in the two ESPs samples.

### Hydrolysis of host proteins by SjCE2b

Recombinant SjCE2b (100 nM) was incubated at 37 °C for 18 h with 20-100 μg of various host proteins, including collagen, elastin, keratin, fibronectin, immunoglobulins (IgA, IgG, and IgM), complement C3, albumin, and hemoglobin (Sigma-Aldrich). The reactions were performed in 50 mM Tris-HCl (pH 8.0) in a total volume of 100 μL. Following incubation, 40 μL of each reaction mixture was resolved by SDS-PAGE using a 4-20% Nupage gel (GenScript) and visualized with Coomassie Brilliant Blue G-250 staining. As a negative control, each protein substrate was incubated without rSjCE2b and processed in parallel under identical conditions.

### Stable isotope dimethyl labeling proteomic analysis of cultivated human epidermis digested with SjCE2b

Proteomic analysis of cultivated human epidermis digested by SjCE2b was performed as previously described (24). Briefly, cultivated human dermis was cut into pieces and placed in a 1.5 mL centrifuge tubes. A 200 μL volume of digestion buffer (50 mM Tris-HCl, pH 8.0) containing 1 μM of either active or inhibited SjCE2b was added. Reactions were mixed by pipetting and incubated for 6 h at 37°C. The inhibited SjCE2b control was prepared by pre-incubating 1 μM SjCE2b with 10 μM MeoSuc-AAPF-CMK for 1 h at room temperature, which completely abolished enzymatic activity against the Suc-AAPF-AMC substrate. After incubation, the reactions were centrifuged at 13,000 × g for 15 min at 4 °C. Finally, a 100 μL aliquot of the supernatant was collected and lyophilized for subsequent mass spectrometry analysis. Downstream sample processing and data analysis were conducted exactly as described previously (24).

### Inhibition of cercariae invasion

Fifteen C57BL/6 mice were randomly divided into three groups (n = 5 per group), an anti-rSjCE2b IgG treated group, a pre-immune IgG treated group, and a PBS-treated control group. Both antibody preparations were diluted in PBS. The abdomens of the mice were shaved prior to exposure. For the anti-rSjCE2b IgG-treated group, 40 cercariae were placed into a 10 mg/mL anti-rSjCE2b IgG solution on a coverslip. Following a 10-min incubation, the coverslip was applied directly to the shaved abdomen of each mouse, with the cercariae-containing side facing down. Parallel control groups were challenged identically using cercariae pre-incubated with either 10 mg/mL pre-immune IgG or PBS. To evaluate invasion efficiency, adult worms were collected and counted 28 days post-infection.

## Results

### Proteomic analysis of *S. japonicum* cercarial secretions induced by linoleic acid or host skin stimulation

Cercarial excretory/secretory products (ESPs) were collected following induction by either chemical stimulation with linoleic acid (Fig 1A) or biological stimulation with mouse skin (Fig 1B). Mass spectrometric analysis identified 74 proteins in the ESPs obtained after mouse skin stimulation (S1 Table) and 84 proteins in those obtained after linoleic acid stimulation (S2 Table). Among these, 49 proteins were unique to the skin-stimulated samples, 59 were unique to the linoleic acid-stimulated samples, and 25 were detected in both groups (Fig 1C). The shared proteins included enzymes involved in glycolysis (pyruvate kinase, glyceraldehyde-3-phosphate dehydrogenase, triosephosphate isomerase, glucose-6-phosphate isomerase, enolase, and fructose-bisphosphate aldolase); proteins associated with antioxidant defense, redox regulation, and stress responses (glutathione S-transferase, heat shock 70 kDa protein, and endoplasmic reticulum chaperone BiP); immunomodulatory proteins (venom allergen-like 4, antigen Sm21.7, and tegument antigen); and proteins involved in calcium signaling (calcium-binding protein, 20 kDa calcium-binding protein, EF-hand domain-containing calcium-binding protein, and 14-3-3 protein epsilon) (S1 and S2 Table).

**Fig 1.**
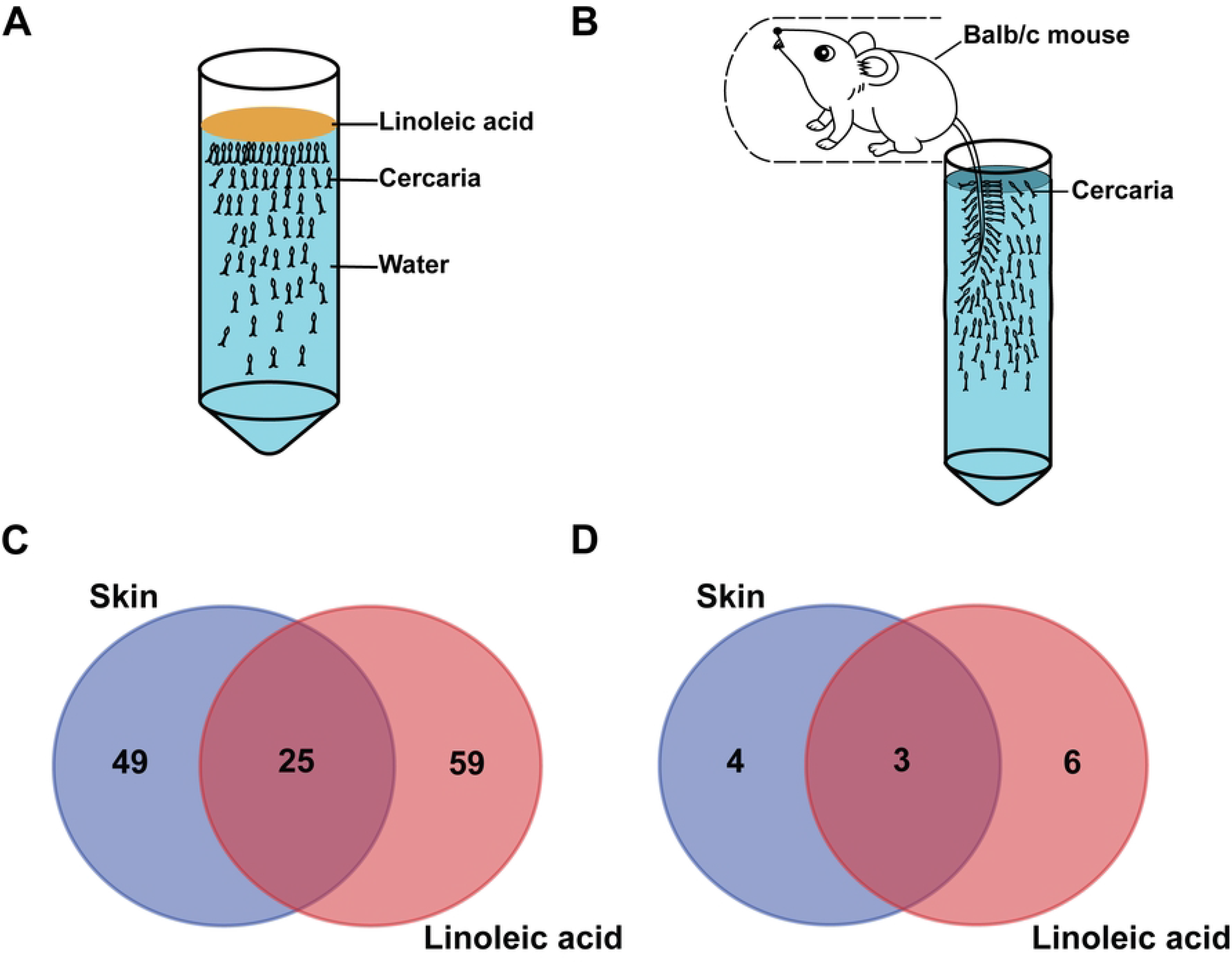
Proteomic analysis of *S. japonicum* cercarial ESPs stimulated by the linoleic acid and host skin. (A) Collection of cercarial ESPs stimulated by linoleic acid. (B) Collection of cercarial ESPs stimulated my mouse skin. (C) Number of proteins identified using the two different stimulation methods. (D). Number of peptidases identified using the two different stimulation methods.

BLASTp searches against the MEROPS 12.5 database identified seven peptidases in the skin-stimulated samples and nine in the linoleic acid-stimulated samples, three of which were shared between the two groups (Fig 1D; Table 1). Notably, while both stimuli successfully triggered the secretion of cercarial elastase (SjCE2b) (S1 Fig), members of the leishmanolysin metalloproteinase family were detected only in the skin-stimulated samples.

### SjCE2b is a secreted trypsin-like serine protease

The SjCE2b open reading frame (ORF) comprises 792 bp and encodes a 264-amino-acid protein with a predicted molecular mass of 28.5 kDa. Sequence homology analysis indicated that SjCE2b contains two major regions: an N-terminal signal peptide domain (residues 1-24) and a Tryp_SPc domain (residues 26-259), which belongs to the trypsin family (Fig 2A). To establish a structural framework for evaluating structure-activity relationships, a three-dimensional model of SjCE2b was generated by homology modelling (Fig 2B). Molecular docking analysis suggested that the inhibitor Suc-AAPF-CMK can access the catalytic pocket and interact with the active site residues of the enzyme (Fig 2C).

**Fig 2.**
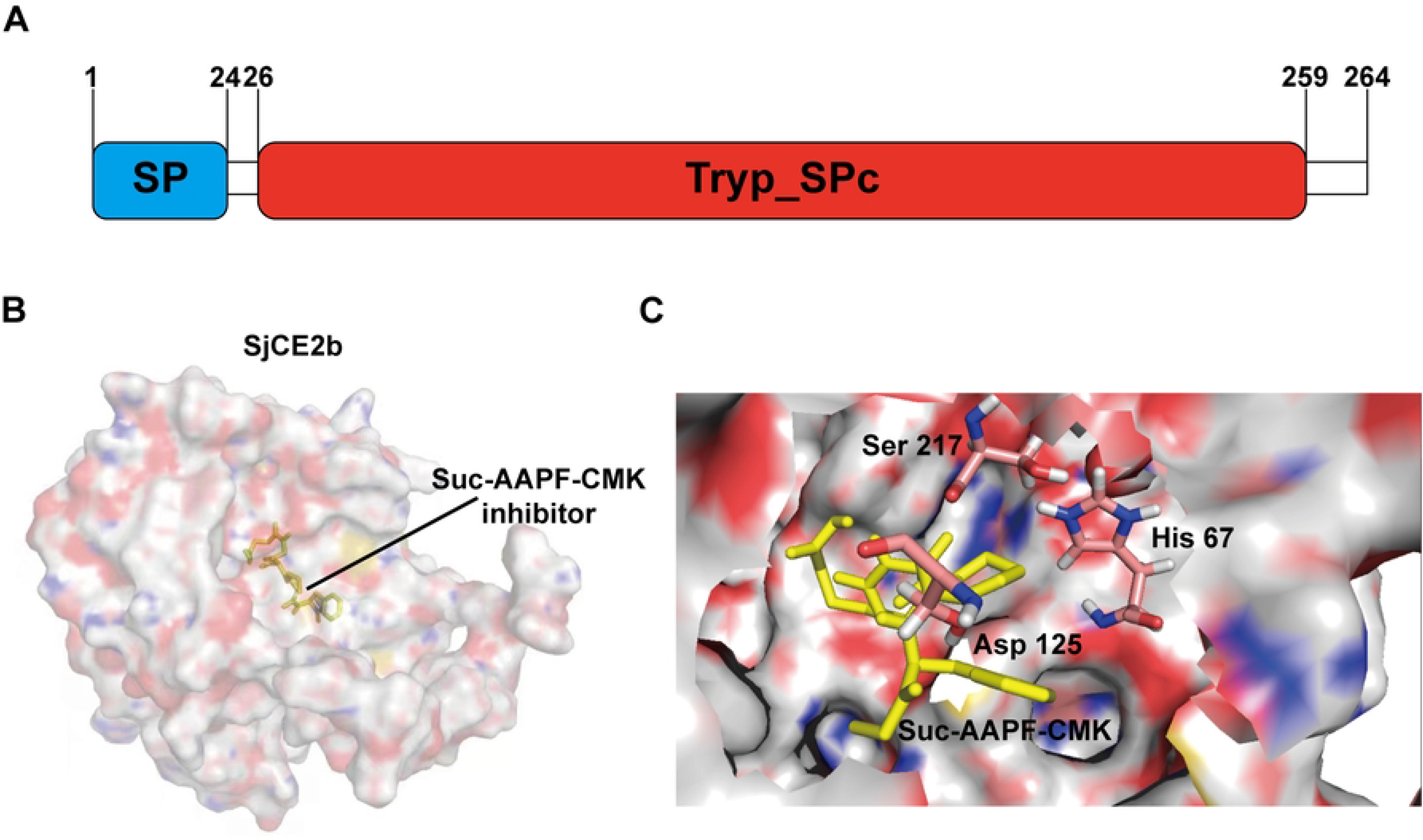
Domain architecture and homology model of SjCE2b. (A) Schematic representation of the domain structure, highlighting the N-terminal signal peptide (SP; blue) and the catalytic Tryp_SPc domain (red). (B) Surface representation of the SjCE2b model in complex with the inhibitor Suc-AAPF-CMK. (C) Close-up view of the interactions between the SjCE2b active site and the inhibitor Suc-AAPF-CMK.

### SjCE2b is highly expressed in cercariae and localizes to the acetabular glands and their ducts

RT-qPCR analysis showed that SjCE2b mRNA was detected only at the sporocyst stage (Fig 3A), western-blot analysis indicated that SjCE2b protein was detected only at the cercaria stage (Fig 3B). Indirect immunofluorescence imaging of fixed cercariae demonstrated that SjCE2b was specifically localized to the cercarial acetabular glands and was extended along their secretory ducts. Notably, intense fluorescence signals concentrated at the distal termini of the ducts, suggesting that the protease is packaged and staged for immediate release during skin penetration (Fig 3C). No signal was detected in the pre-immune IgG or PBS controls (Fig 3C).

**Fig 3.**
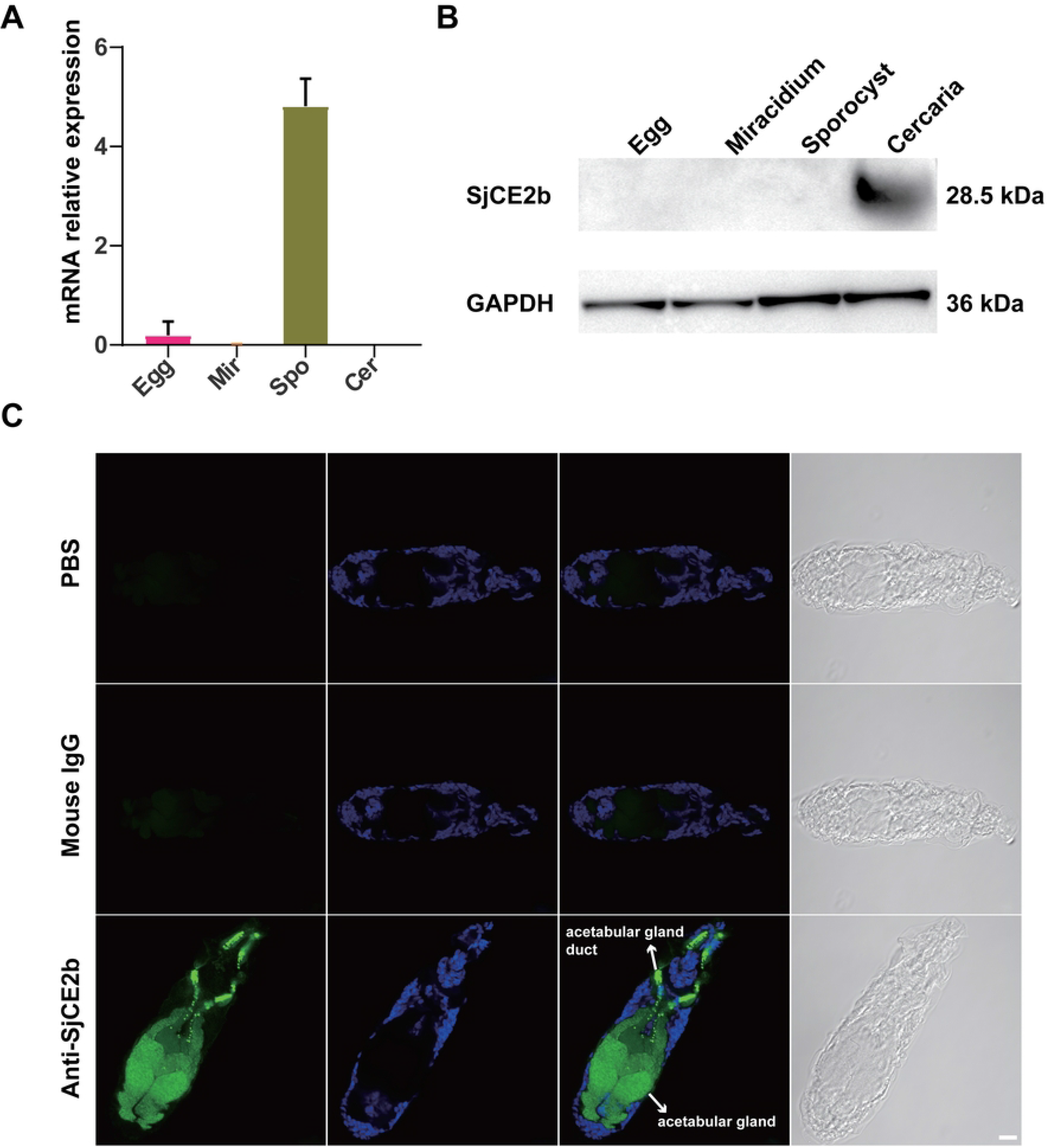
Developmental expression profiles and localization of SjCE2b. (A) Relative SjCE2b mRNA abundance across four larval stages quantified via RT-qPCR. Egg, egg; Mir, miracidium; Spo, sporocyst; Cer, cercaria. (B) Stage-specific protein expression of SjCE2b assessed by western-blot, with GAPDH served as an internal control. (C) Immunofluorescence imaging showing SjCE2b localization in the acetabular glands and secretory ducts in cercariae, indicated by white arrows. SjCE2b signals are shown in green. Scale bar, 10 μm.

### Enzymatic activity of SjCE2b

Recombinant SjCE2b was successfully expressed in the yeast *Pichia Pastoris* (S2A Fig). Western-blot analysis using an anti-rSjCE2b IgG of the purified protein revealed an apparent molecular mass of approximately 30 kDa, aligning closely with the predicted theoretical size of 28.5 kDa (S2B Fig). The enzymatic activity of SjCE2b was assessed using the fluorogenic substrate Suc-AAPF-AMC. A dose-dependent assay showed that substrate degradation increased with increasing enzyme concentration (Fig 4A). The inhibitor specificity of SjCE2b was further examined using a panel of small molecule and protein inhibitors, as listed in Table 2. SjCE2b activity was completely inhibited by the selective elastase inhibitor MeoSuc-AAPF-CMK. Partial inhibition was observed with leupeptin, E-64, pepstatin A, and benzamidine-inhibitors targeting serine and cysteine protease, cysteine protease, aspartic proteases, and trypsin-like serine proteases, respectively (Table 2).

**Fig 4.**
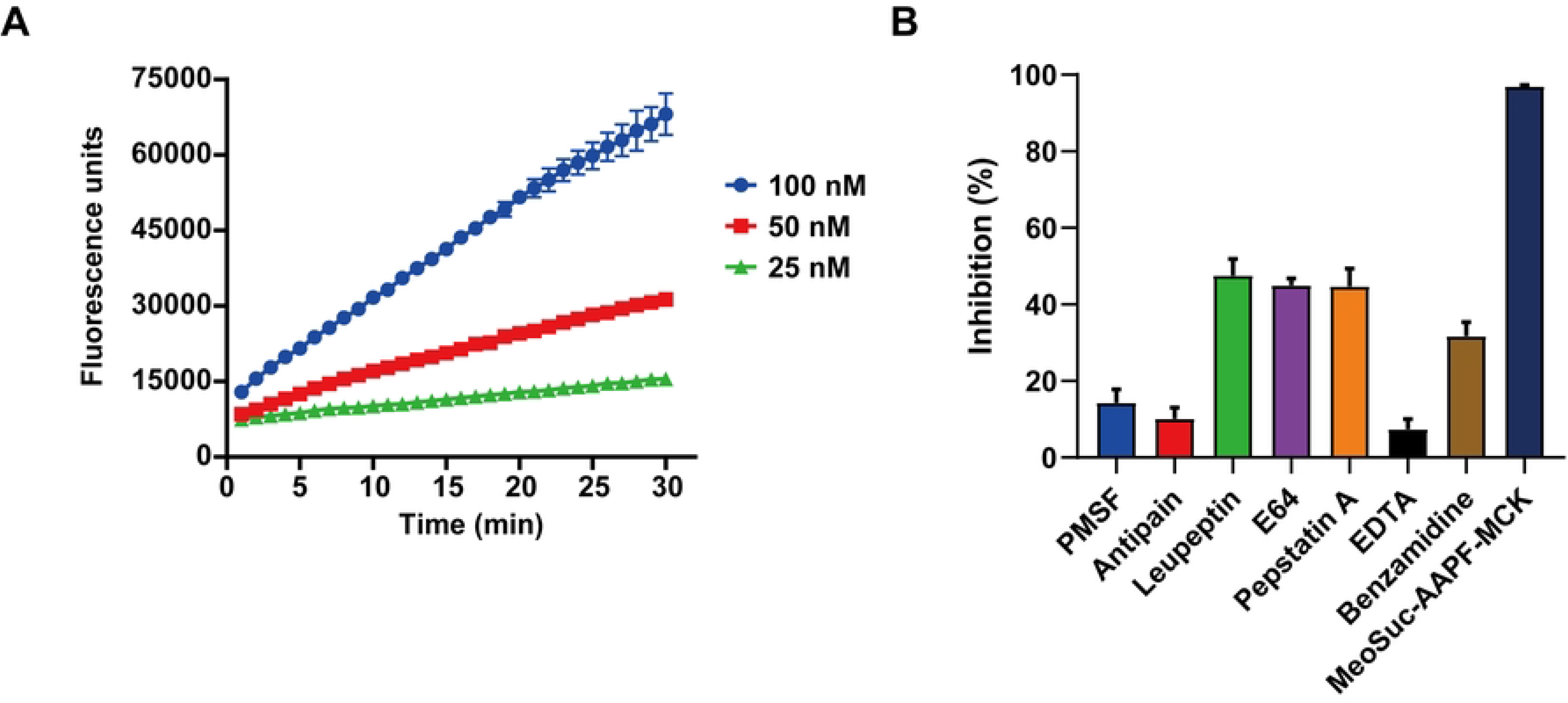
Enzymatic activity and inhibition profile of recombinant SjCE2b. (A) Dose-dependent enzymatic activity of SjCE2b measured by cleavage of the fluorogenic substrate Suc-AAPF-AMC. Data are represented as the mean ± S.D. of three replicates. (B) Inhibition of recombinant SjCE2b by protease inhibitors. The enzyme was pre-incubated with each inhibitor, and residual activity was measured in a kinetic assay using the fluorogenic substrate Suc-AAPF-AMC. Mean values ± S.D. from three replicates are expressed as percentage inhibition relative to the uninhibited control.

**Table 2.** Inhibition of recombinant SjCE2b by protease inhibitors.

| Inhibitor<sup>a</sup> | Target protease<sup>b</sup> | Concentration (μM) | Inhibition (%) |
| --- | --- | --- | --- |
| PMSF | SP | 1000 | 14.3±3.7 |
| Antipain | SP, CP | 20 | 10.1±3.0 |
| Leupeptin | SP, CP | 20 | 47.5±4.4 |
| E64 | CP | 10 | 45.0±1.8 |
| Pepstatin A | AP | 1 | 44.7±4.7 |
| EDTA | MP | 1000 | 7.3±2.7 |
| Benzamidine | SP (trypsin type) | 10 | 31.7±3.7 |
| MeoSuc-AAPF-CMK | SP (elastase) | 10 | 97.0±0.3 |

<sup>a</sup>Abbreviations: PMSF, phenylmethylsulfonyl fluoride; E64, trans-Epoxy succinyl-L-leucylamido(4-guanidino) butane; EDTA, ethylenediaminetetraacetic acid.
<sup>b</sup>Target proteases are classified according to catalytic type as aspartic protease (AP),
cysteine protease (CP), serine protease (SP) and metalloprotease (MP).

### Hydrolysis of host proteins

To elucidate the potential functional roles of SjCE2b during cercariae invasion, its proteolytic activity was evaluated against a panel of host macromolecules, including skin-associated proteins (collagen, elastin, keratin, and fibronectin), immune system components (immunoglobulin A, immunoglobulin G, immunoglobulin M and complement C3), and blood components (albumin and hemoglobin). SjCE2b efficiently degraded collagen, keratin, fibronectin and complement C3. Elastin, immunoglobulin A, and immunoglobulin G were partially digested. No hydrolysis was observed for hemoglobin or serum albumin, two major protein components of vertebrate host blood (Fig 5).

**Fig 5.**
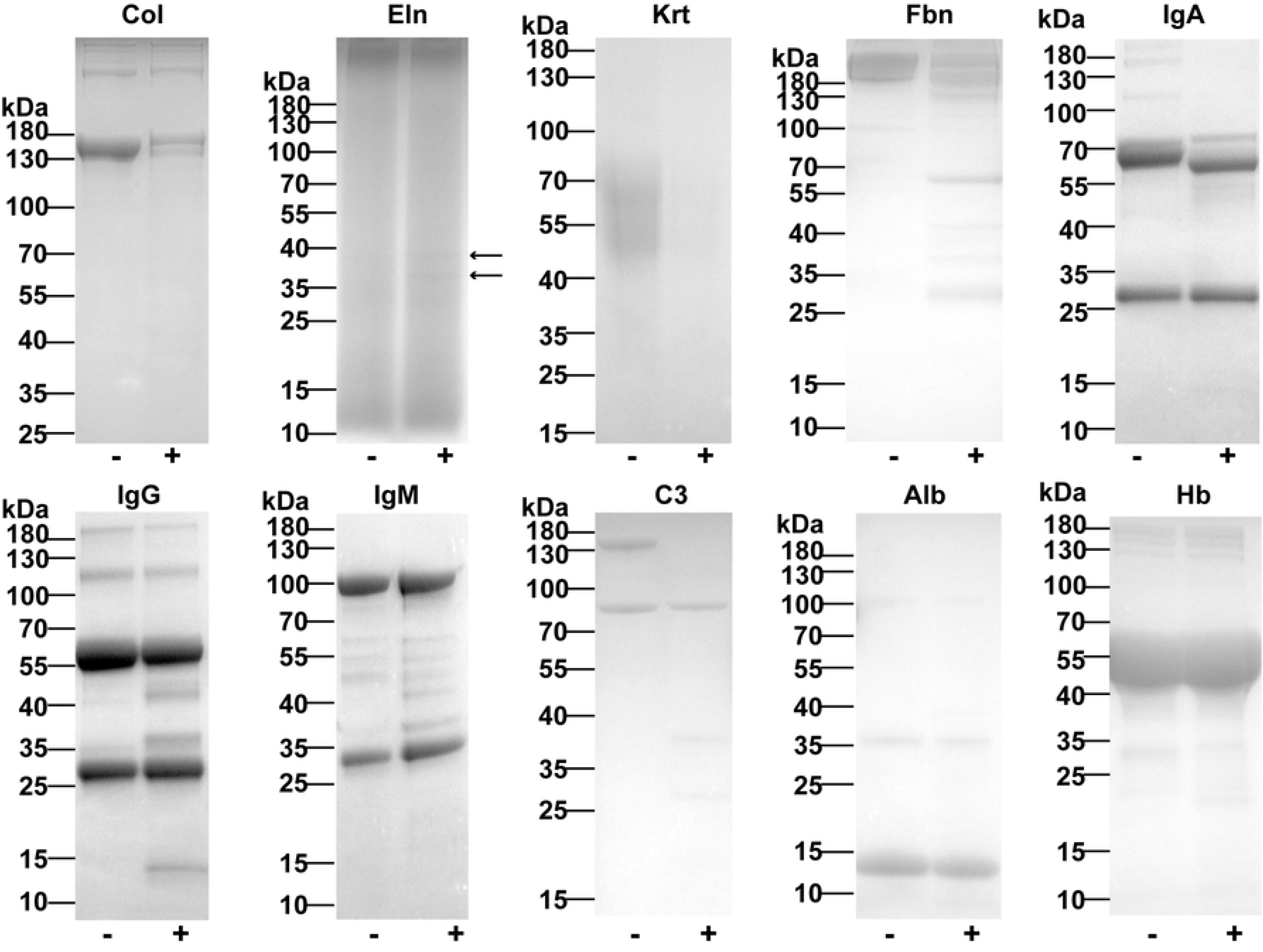
Digestion of selected natural substrates by recombinant SjCE2b. Proteolytic activity of recombinant SjCE2b was assessed against major host skin components, including collagen (Col), elastin (Eln), keratin (Krt) and fibronectin (Fbn); immune system components, including immunoglobulin A (IgA), immunoglobulin G (IgG), immunoglobulin M (IgM) and complement C3 (C3); and blood components, including albumin (Alb) and hemoglobin (Hb). Arrows in the Eln panel indicate small fragments generated by SjCE2b digestion. Substrates were incubated in 50 mM Tris HCl buffer (pH 8.0) in the presence (+) or absence (-) of SjCE2b for 18 h at 37°C and analyzed by 10% SDS-PAGE. Molecular weight markers of each substrate are shown on the left side.

### Hydrolysis of human epidermal proteins by SjCE2b

A total of 58 human epidermal proteins were identified as proteolytic substrates of SjCE2b (Table 3). These include one cornified layer protein, periplakin; four granular layer proteins, filaggrin, filaggrin-2, dermokine and caspase-14; ten keratins; and seven cytoskeletal proteins, including myosin-9, desmoplakin, plakophilin-3, actin (cytoplasmic 1), actin (cytoplasmic 2), protein enabled homolog and PDZ and LIM domain protein 1. In addition, eight extracellular proteins, including serpin A12, annexin A1, interleukin-36, and alpha-2-macroglobulin-like protein 1, were also cleaved by SjCE2b.

**Table 3.**
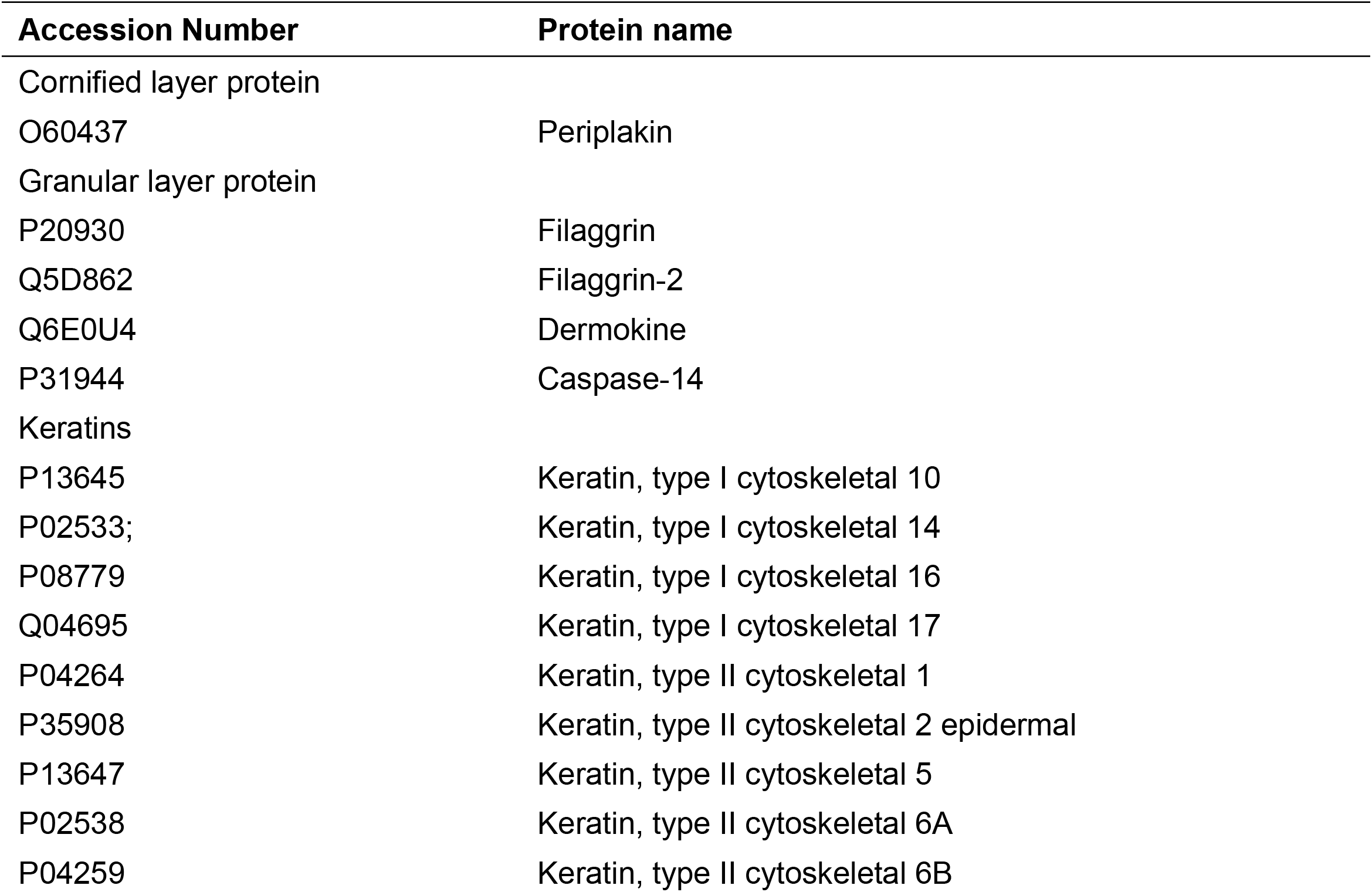
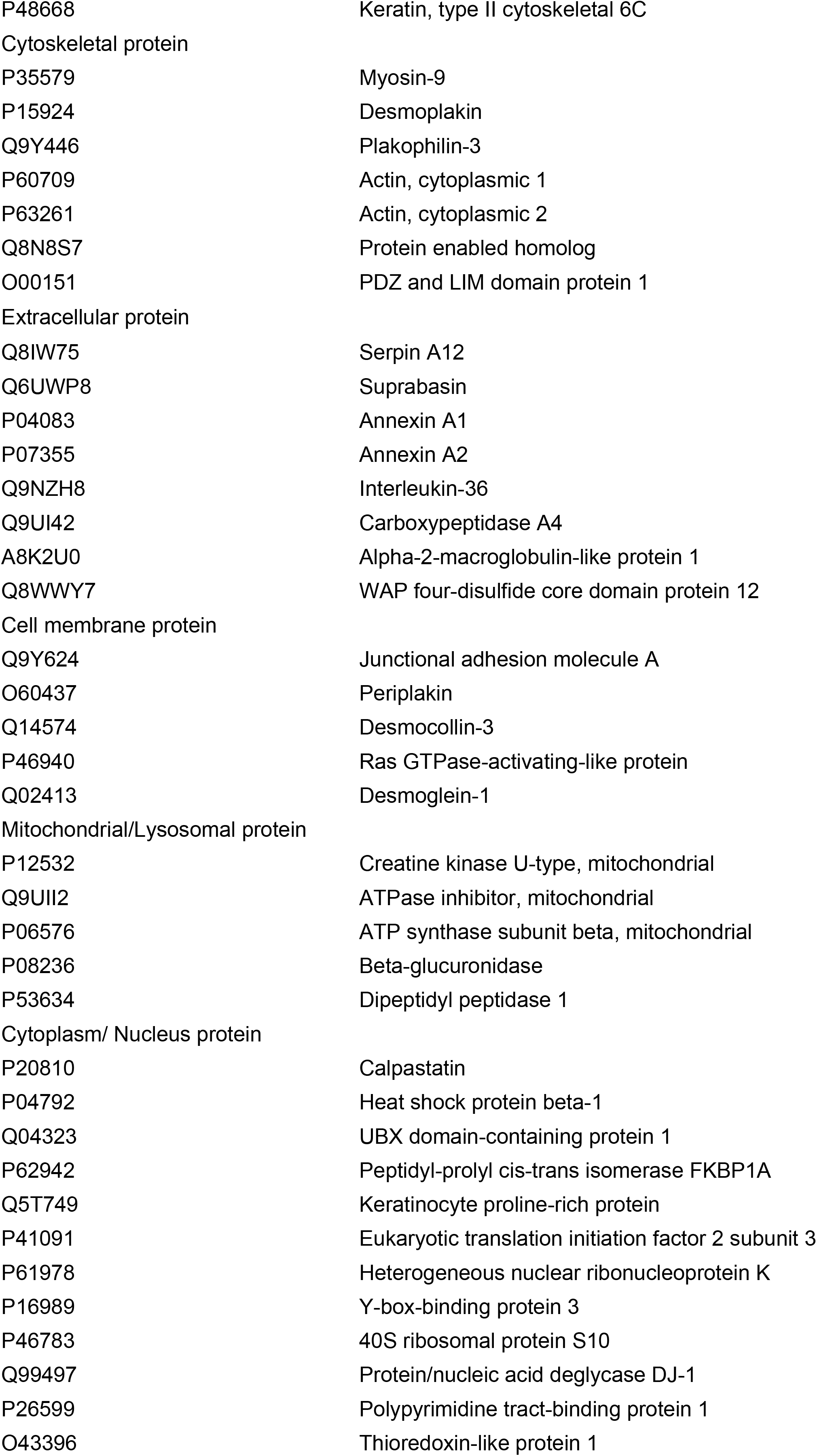
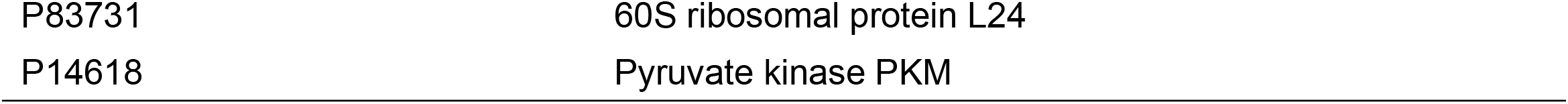
Substrates of SjCE2b detected in cultivated human epidermis.

### Inhibition of parasite invasion by antibody against SjCE2b

Invasion inhibition assays were performed to evaluate the contribution of SjCE2b in *S. japonicum* skin penetration. Compared with the pre-immune IgG control group, the anti-rSjCE2b-treated group showed a significant reduction in worm burden (80.85%, *P*-value < 0.05). No significant difference was observed between the PBS control group and the pre-immune IgG control group (Fig 6).

**Fig 6.**
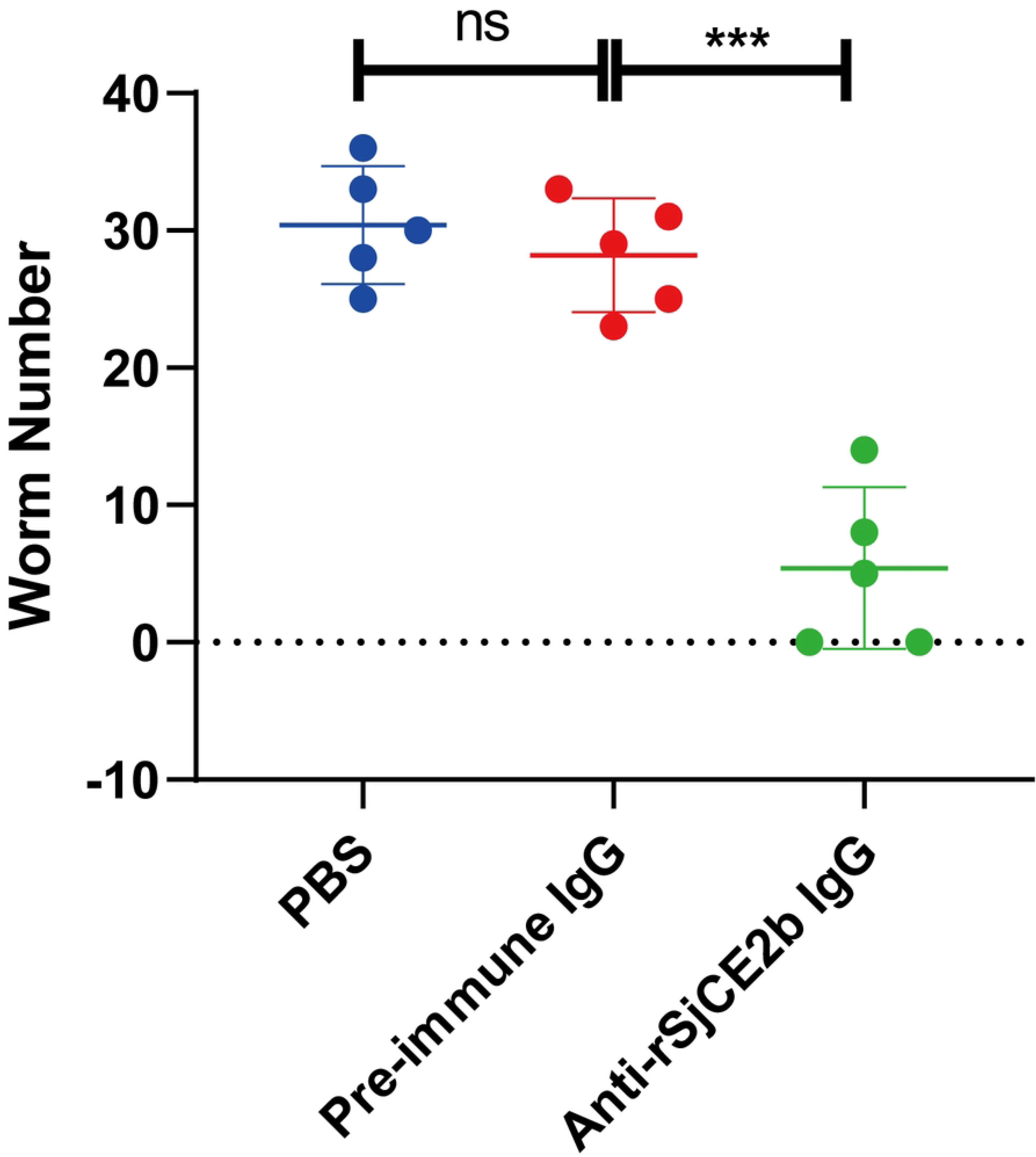
Inhibition of cercarial invasion by anti-rSjCE2b IgG. Mice were experimentally challenged with 40 *S. japonicum* cercariae, and adult worms were collected 28 days post infection. Statistically significant difference (*** = p-value < 0.001).

## Discussion

Skin penetration represents the critical first step in definitive host invasion by *Schistosoma* species. During this process, ESPs, particularly proteases released from the acetabular glands of infective cercariae, degrade host skin components to facilitate parasite entry and transmission. While SmCE is well established as a primary driver of tissue penetration (10), the functional contribution of SjCE2b during *S. japonicum* cercariae invasion remains unclear, largely due to the lack of a soluble, catalytically active recombinant enzyme. In this study, we demonstrate that the native SjCE2b is secreted by cercariae in response to linoleic acid or direct host skin contact. Furthermore, we generated enzymatically active recombinant SjCE2b in *P. pastoris* and experimentally confirmed its essential role in host skin penetration.

We performed the first comprehensive proteomic characterization of the *S. japonicum* cercarial ESPs induced by either linoleic acid or host skin. While both stimuli consistently triggered the secretion of SjCE2b, three distinct leishmanolysins were identified only in the skin-stimulated samples. Leishmanolysins (also known as invadolysins) play important roles in cercariae invasion; for instance, *S. mansoni* cercarial invadolysin 1 (SmCI-1) was identified as the second most abundant protein released from cercariae acetabular glands during transformation to schistosomula (7). Functionally, SmCI-1 promotes larval survival within host skin by degrading extracellular matrix components and inactivating complement C3b (25). Previously, we demonstrated that *S. japonicum* leishmanolysin-like peptidase isoform 1 (SjLLPi1) facilitates definitive host penetration by hydrolyzing skin components (26).

An unexpected finding of our proteomic data was the absence of detectable cathepsins. Dvořák et al., who detected cathepsin B-but not elastase—in *in vitro*-collected *S. japonicum* secretions, reporting ∼40-fold higher cathepsin B-like activity relative to *S. mansoni* (6). Because we previously localized the cathepsin B-SjCB2 to cercarial acetabular glands and confirmed its capacity to digest both skin and immune proteins (24), we postulate that SjCB2 is stored at low abundance within the glands yet possesses high specific enzymatic activity. Consequently, future detection of SjCB2 in cercarial ESPs will likely require advanced mass spectrometry platforms, such as the Orbitrap Astral, which offer the requisite sensitivity to resolve low-abundance proteins (27). Supporting this hypothesis, cathepsins were similarly absent from *S. mansoni* secretomes induced via skin lipids or mechanical transformation (7, 8). In addition to proteases, several calcium-binding proteins and venom allergen-like proteins were identified in the ESPs of both stimuli. Calcium-binding proteins, such as SjCa8 (28), have been implicated in immune evasion by cercariae, whereas venom allergen-like (VAL) proteins have been reported to bind lipids, as shown for SmVAL4 (29), or plasminogen, as shown for SmVAL18 (30).

SjCE2b mRNA was detected only at the sporocyst stage, whereas the corresponding protein was restricted to the cercariae stage. Previous studies have shown that mature *Schistosoma* cercariae are transcriptionally silent-a repressive chromatin state maintained by the histone mark H3K27me3 (31), and exhibit limited translational activity within the cercarial head (32). We therefore speculate that SjCE2b may be transcribed in the sporocyst and translated subsequently during cercariae development. Similar expression patterns have been reported for SjLLPi1 (26). Immunofluorescence microscopy revealed that SjCE2b localized to the acetabular glands, with dense accumulation along the secretory ducts. Moreover, the rapid clearance of SjCE2b upon transformation into schistosomula (33) indicates that this protease functions specifically in the skin penetration process.

Overcoming prior limitations where *Escherichia coli* expression yielded insoluble inclusion bodies (34), we successfully generated soluble, recombinant SjCE2b in *P. pastoris*. The purified enzyme exhibited dose-dependent proteolytic cleavage of the fluorogenic substrate Suc-AAPF-AMC; this activity was completely abolished by the selective inhibitor MeoSuc-AAPF-CMK, confirming that SjCE2b functions as a trypsin-like serine protease. Given its rapid consumption during host invasion, SjCE2b likely plays a pivotal role in tissue degradation. Proteomic profiling revealed that SjCE2b cleaved numerous human epidermal proteins, including essential stratum corneum barrier components such as filaggrin (35) and filaggrin-2 (36). In addition, the enzyme degraded ten distinct type I and type II keratins, indicating direct disruption of the keratin intermediate filaments that confer mechanical resilience to epithelial tissues (37). Furthermore, SjCE2b cleaved major desmosomal constituents—including desmoglein-1, desmocollin-3, desmoplakin, and plakophilin-3—thereby targeting the intercellular junctions that anchor keratin networks between adjacent keratinocytes (38). Together, these findings suggest that SjCE2b facilitates *S. japonicum* cercarial penetration through the epidermis by progressively weakening the cornified envelope, degrading keratin-based mechanical scaffolds, and disrupting epidermal cell–cell adhesion.

Using an *in vitro* digestion assay, we found that SjCE2b was capable of degrading collagen and elastin—the primary structural scaffolds of the dermis—as well as the glycoprotein fibronectin, which provides essential cellular adhesion cues (39). These proteolytic activities indicate that SjCE2b can disrupt the physical structure of the host dermis. In addition, SjCE2b cleaved complement C3 and immunoglobulins (IgA and IgG), suggesting that it may facilitate evasion of the dermal immune response through proteolysis of immunoglobins and complement factors. Similar complement cleavage activity has been reported for SmCE (14). Finally, to directly evaluate the contribution of SjCE2b to host invasion, we neutralized the secreted native enzyme using anti-rSjCE2b IgG. This treatment resulted in an 80.85% reduction in adult worm burden— a substantially greater attenuation than previously observed for SjCB2 (22.94%) (24), thus firmly establishing SjCE2b as a principal mediator of *S. japonicum* skin penetration.

In summary, our findings indicate that SjCE2b is secreted from the acetabular glands upon contact with host skin. This protease facilitates the degradation of structural proteins within the epidermis and extracellular matrix (ECM) components in the dermis, enabling the parasite to breach the epidermal-dermal barrier and gain access to the vasculature or lymphatic system. Additionally, SjCE2b may promote the evasion of host dermal immune responses through the cleavage of complement factors and immunoglobulins. Ultimately, the profound impact of neutralizing this enzyme—which yielded an 80.85% reduction in adult worm burden—firmly establishes SjCE2b as a compelling therapeutic target for the development of novel pharmacological interventions or transmission-blocking vaccines against *S. japonicum*.

## Acknowledgment

We gratefully acknowledge Ying Wu, Xiaojin Mo, Bin Xu, and Yang Hong from the National Institute of Parasitic Diseases, Chinese Center for Disease Control and Prevention, for their sustained support and provision of parasite materials. We also thank Shanghai ReMed Biotechnology Co., Ltd. for supplying the cultivated human epidermal tissues.

## Supporting Information

S1 Fig. Identification of SjCE2b in *S. japonicum* cercarial ESPs under two stimulation conditions. (A) Two unique peptides corresponding to SjCE2b were identified in the respective ESP preparations. (B) Identification of the peptide TIGNPICIQPGPDEK in linoleic acid-stimulated ESPs samples. (C) Identification of the peptide AVNHDASEIKIPPEYEPTCQLK in host skin-stimulated ESPs samples.

S2 Fig. Expression and purification of recombinant SjCE2b in *P. pastoris*. (A) Coomassie blue-stained 10% SDS–PAGE gel of purified SjCE2b. (B) Western blot analysis of purified SjCE2b using an anti-rSjCE2b polyclonal antibody.

S1 Table. Proteins identified by proteomic analysis of mouse skin-induced ESPs samples.

S2 Table. Proteins identified by proteomic analysis of linoleic acid-induced ESPs samples.

## References

1. Buonfrate D, Ferrari TCA, Adegnika AA, Russell Stothard J, Gobbi FG. Human schistosomiasis. The Lancet. 2025;405(10479):658–70.

2. McManus DP, Dunne DW, Sacko M, Utzinger J, Vennervald BJ, Zhou XN. Schistosomiasis. Nat Rev Dis Primers. 2018;4(1):13.

3. McKerrow JH, Salter J. Invasion of skin by *Schistosoma* cercariae. Trends Parasitol. 2002;18(5):193–5.

4. Panzner U, Utzinger J, Keiser J. Schistosomiasis: cercarial finding and recognizing of human hosts as a prerequisite of invasion. Clin Microbiol Rev. 2025;38(3):e0019624.

5. Hambrook JR, Hanington PC. Immune Evasion Strategies of Schistosomes. Front Immunol. 2020;11:624178.

6. Dvorák J, Mashiyama ST, Braschi S, Sajid M, Knudsen GM, Hansell E, et al. Differential use of protease families for invasion by schistosome cercariae. Biochimie. 2008;90(2):345–58.

7. Curwen RS, Ashton PD, Sundaralingam S, Wilson RA. Identification of novel proteases and immunomodulators in the secretions of schistosome cercariae that facilitate host entry. Mol Cell Proteomics. 2006;5(5):835–44.

8. Knudsen GM, Medzihradszky KF, Lim KC, Hansell E, McKerrow JH. Proteomic analysis of *Schistosoma mansoni* cercarial secretions. Mol Cell Proteomics. 2005;4(12):1862–75.

9. Hansell E, Braschi S, Medzihradszky KF, Sajid M, Debnath M, Ingram J, et al. Proteomic analysis of skin invasion by blood fluke larvae. PLoS Negl Trop Dis. 2008;2(7):e262.

10. Ingram JR, Rafi SB, Eroy-Reveles AA, Ray M, Lambeth L, Hsieh I, et al. Investigation of the proteolytic functions of an expanded cercarial elastase gene family in *Schistosoma mansoni*. PLoS Negl Trop Dis. 2012;6(4):e1589.

11. Young ND, Jex AR, Li B, Liu S, Yang L, Xiong Z, et al. Whole-genome sequence of *Schistosoma haematobium*. Nat Genet. 2012;44(2):221–5.

12. The *Schistosoma japonicum* genome reveals features of host-parasite interplay. Nature. 2009; 460(7253):345–51.

13. Salter JP, Lim KC, Hansell E, Hsieh I, McKerrow JH. Schistosome invasion of human skin and degradation of dermal elastin are mediated by a single serine protease. J Biol Chem. 2000;275(49):38667–73.

14. Ingram J, Knudsen G, Lim KC, Hansell E, Sakanari J, McKerrow J. Proteomic analysis of human skin treated with larval schistosome peptidases reveals distinct invasion strategies among species of blood flukes. PLoS Negl Trop Dis. 2011;5(9):e1337.

15. Lim KC, Sun E, Bahgat M, Bucks D, Guy R, Hinz RS, et al. Blockage of skin invasion by schistosome cercariae by serine protease inhibitors. Am J Trop Med Hyg. 1999;60(3):487–92.

16. Shiff CJ, Cmelik SH, Ley HE, Kriel RL. The influence of human skin lipids on the cercarial penetration responses of *Schistosoma haematobium* and *Schistosoma mansoni*. J Parasitol. 1972;58(3):476–80.

17. Wiœniewski JR, Zougman A, Nagaraj N, Mann M. Universal sample preparation method for proteome analysis. Nat Methods. 2009;6(5):359–62.

18. Kovalchuk SI, Jensen ON, Rogowska-Wrzesinska A. FlashPack: Fast and Simple Preparation of Ultrahigh-performance Capillary Columns for LC-MS. Mol Cell Proteomics. 2019;18(2):383–90.

19. Luo F, Yang W, Yin M, Mo X, Pang Y, Sun C, et al. A chromosome-level genome of the human blood fluke *Schistosoma japonicum* identifies the genomic basis of host-switching. Cell Rep. 2022;39(1):110638.

20. Rawlings ND, Barrett AJ, Thomas PD, Huang X, Bateman A, Finn RD. The MEROPS database of proteolytic enzymes, their substrates and inhibitors in 2017 and a comparison with peptidases in the PANTHER database. Nucleic Acids Res. 2018;46(D1):D624–d32.

21. Leontovyè A, Ulrychová L, O’Donoghue AJ, Vondrášek J, Marešová L, Hubálek M, et al. SmSP2: A serine protease secreted by the blood fluke pathogen *Schistosoma mansoni* with anti-hemostatic properties. PLoS Negl Trop Dis. 2018;12(4):e0006446.

22. Fajtová P, Štefaniæ S, Hradilek M, Dvoøák J, Vondrášek J, Jílková A, et al. Prolyl Oligopeptidase from the Blood Fluke *Schistosoma mansoni*: From Functional Analysis to Anti-schistosomal Inhibitors. PLoS Negl Trop Dis. 2015;9(6):e0003827.

23. Livak KJ, Schmittgen TD. Analysis of relative gene expression data using real-time quantitative PCR and the 2(-Delta Delta C(T)) Method. Methods. 2001;25(4):402–8.

24. Zhu B, Luo F, Shen Y, Yang W, Sun C, Wang J, et al. *Schistosoma japonicum* cathepsin B2 (SjCB2) facilitates parasite invasion through the skin. PLoS Negl Trop Dis. 2020;14(10):e0008810.

25. Hambrook JR, Hanington PC. A cercarial invadolysin interferes with the host immune response and facilitates infection establishment of *Schistosoma mansoni*. PLoS Pathog. 2023;19(2):e1010884.

26. Chen F, Zhu B, Fang Y, Li Z, Lei Z, Xue Z, et al. *Schistosoma japonicum* leishmanolysin SjLLPi1 facilitates the invasion of cercariae into the host skin. PLoS Pathog. 2025;21(8):e1013446.

27. Hendricks NG, Bhosale SD, Keoseyan AJ, Ortiz J, Stotland A, Seyedmohammad S, et al. An Inflection Point in High-Throughput Proteomics with Orbitrap Astral: Analysis of Biofluids, Cells, and Tissues. J Proteome Res. 2024;23(9):4163–9.

28. Liu J, Pan T, You X, Xu Y, Liang J, Limpanont Y, et al. SjCa8, a calcium-binding protein from *Schistosoma japonicum*, inhibits cell migration and suppresses nitric oxide release of RAW264.7 macrophages. Parasit Vectors. 2015;8:513.

29. Kelleher A, Darwiche R, Rezende WC, Farias LP, Leite LC, Schneiter R, et al. *Schistosoma mansoni* venom allergen-like protein 4 (SmVAL4) is a novel lipid-binding SCP/TAPS protein that lacks the prototypical CAP motifs. Corrigendum. Acta Crystallogr D Biol Crystallogr. 2015;71(Pt 4):1022.

30. Fernandes RS, Fernandes LGV, de Godoy AS, Miyasato PA, Nakano E, Farias LP, et al. *Schistosoma mansoni* venom allergen-like protein 18 (SmVAL18) is a plasminogen-binding protein secreted during the early stages of mammalian-host infection. Mol Biochem Parasitol. 2018;221:23–31.

31. Roquis D, Lepesant JM, Picard MA, Freitag M, Parrinello H, Groth M, et al. The Epigenome of *Schistosoma mansoni* Provides Insight about How Cercariae Poise Transcription until Infection. PLoS Negl Trop Dis. 2015;9(8):e0003853.

32. Hagerty JR, Jolly ER. Heads or tails? Differential translational regulation in cercarial heads and tails of schistosome worms. PLoS One. 2019;14(10):e0224358.

33. Liu M, Ju C, Du XF, Shen HM, Wang JP, Li J, et al. Proteomic Analysis on Cercariae and Schistosomula in Reference to Potential Proteases Involved in Host Invasion of *Schistosoma japonicum* Larvae. J Proteome Res. 2015;14(11):4623–34.

34. Zhang T, Mo XJ, Xu B, Yang Z, Gobert GN, Yan S, et al. Enzyme activity of *Schistosoma japonicum* cercarial elastase SjCE-2b ascertained by in vitro refolded recombinant protein. Acta Trop. 2018;187:15–22.

35. Levin J, Friedlander SF, Del Rosso JQ. Atopic dermatitis and the stratum corneum: part 1: the role of filaggrin in the stratum corneum barrier and atopic skin. J Clin Aesthet Dermatol. 2013;6(10):16–22.

36. Wang Z, Chen H, Wang Y, Wu C, Ye T, Xia H, et al. Recombinant filaggrin-2 improves skin barrier function and attenuates ultraviolet B (UVB) irradiation-induced epidermal barrier disruption. Int J Biol Macromol. 2024;281(Pt 1):136064.

37. Pora A, Yoon S, Dreissen G, Hoffmann B, Merkel R, Windoffer R, et al. Regulation of keratin network dynamics by the mechanical properties of the environment in migrating cells. Sci Rep. 2020;10(1):4574.

38. Kouklis PD, Hutton E, Fuchs E. Making a connection: direct binding between keratin intermediate filaments and desmosomal proteins. J Cell Biol. 1994;127(4):1049–60.

39. Pfisterer K, Shaw LE, Symmank D, Weninger W. The Extracellular Matrix in Skin Inflammation and Infection. Front Cell Dev Biol. 2021;9:682414.

